# CANCAN: high-resolution copy number and mutation heterogeneity analysis of DNA sequence data for clinical applications

**DOI:** 10.64898/2026.05.16.725235

**Authors:** Arne V Pladsen, Daniel Vodak, Sen Zhao, Sigve Nakken, Daniel Nebdal, Tonje Lien, Britina Kjuul Danielsen, Caroline Wang, Wanja Kildal, Geir Olav Hjortland, Olav Engebråten, Eivind Hovig, Hege G Russnes, Ole Christian Lingjærde

## Abstract

High-throughput DNA sequencing is central to precision oncology, yet robust and interpretable methods for integrated analysis of copy number alterations and somatic variants across sequencing platforms remain limited. We present CANCAN (Copy number integrative ANalysis in CANcer), a platform-agnostic computational framework for high-resolution analysis of allele-specific copy number and variant data. CANCAN integrates novel normalization and segmentation strategies and enables inference of tumor purity, ploidy, subclonality and mutation multiplicity, while providing statistical confidence estimates and transparent evaluation of alternative solutions. Benchmarking across whole-genome, whole-exome and targeted sequencing datasets from TCGA and the IMPRESS-Norway study demonstrates high concordance with established methods, with particularly strong performance on targeted sequencing data. CANCAN accurately estimates global genomic features, including purity and ploidy, even at reduced sequencing coverage, and shows comparable or improved agreement relative to existing tools. In addition, it provides detailed visualization of the genomic context of clinically relevant biomarkers, supporting diagnostic interpretation. CANCAN constitutes a reproducible and interpretable approach for integrated genomic analysis, addressing key methodological and practical challenges in clinical cancer genomics.

## 1 Introduction

High-throughput DNA sequencing is increasingly used to identify biomarkers for predicting treatment response and prognosis in cancer^1,2^. The genomic context of a biomarker has potentially significant clinical implications. For instance, a loss-of-function mutation in a tumor suppressor gene such as *BRCA1* leads to homologous recombination deficiency primarily in the context of biallelic deactivation, which can occur through deletion of the wild-type allele^3,4^. Determining whether the wild-type allele is conserved in the tumor requires a comprehensive analysis that integrates allele-specific absolute copy number, somatic variant data, and the subclonality state^5^. This places high demands on analytical methods. Furthermore, patients may undergo DNA sequencing using various short-read platforms, ranging from targeted panels to whole-genome sequencing. This diversity necessitates presenting genomic results in a way that allows seamless comparison across platforms. Lastly, bioinformatics analyses may occasionally yield ambiguous copy number results^6,7^. In clinical settings, where a single biomarker might determine the treatment trajectory of a patient, it is essential that bioinformatics methods provide transparency regarding such ambiguities, empowering clinicians to make informed decisions.

Although numerous sequencing-based bioinformatics tools exist for analyzing integrated copy number and somatic variant data^7–10^, few are truly platform-agnostic and capable of handling data from a wide range of sequencing approaches — from highly targeted panels to whole-genome sequencing. Furthermore, few methods provide detailed insights into the genomic context of specific biomarkers, or statistical confidence estimates for their results. This highlights the need for more integrated diagnostic pipelines that address critical requirements, including robustness, standardization, platform compatibility, interpretability, and effective data presentation. Moreover, the complete analysis pipeline - from raw sequence reads to high-level statistical outputs - should be fully reproducible across laboratories to enable consistent and distributed analyses in multi-center clinical studies.

We here present Copy number integrative ANalysis in CANcer (CANCAN), a tool for analyzing and presenting copy number and somatic variant data in an integrative manner. CANCAN is specifically designed for use in precision cancer diagnostics, to address the described needs met in a clinical setting. The tool brings together several novel strategies for normalization and feature extractions from copy number data and leverages all available sequence reads to enhance the resolution and robustness of the analysis. CANCAN is compatible with data from a wide range of sequencing platforms currently used in the clinic.

## 2 Methods

### 2.1 Copy number profiles and change points

We assume a fixed reference genome where the chromosomes consist of *l*_1_,..., *l*_24_ nucleotides, respectively. The *j*th nucleotide of the *k*th chromosome is assigned the genomic coordinate *t* = *t*(*j, k*) = *j* + *l*_0_ + ... + *l*_*k*→1_ where *l*_0_ = 0. Possible genomic coordinates are thus *G* = {1, 2,...,*N*} with *N* = *l*_1_ + *…* + *l*_24_. A copy number profile is a mapping *µ* : *G*→ ℝ, and a change point of *µ* is a genomic coordinate t that satisfies *µ*(*t*−1) *≠ µ*(*t*). In the following, we sought to determine from DNA sequence data both the total copy number *µ*(*t*) and its decomposition *µ*(*t*) = *µ*^0^(*t*)+ *µ*^1^(*t*) into allelic copy number profiles.

### 2.2 Total copy ratio

GATK (version 4.2.0.0)^11^ was used to calculate genome-wide total copy ratios *R*_1_,..., *R*_*n*_ for each sample. In brief, each chromosome was subdivided into n non-overlapping bins of width W nucleotides (with a smaller bin at the end of each chromosome). Bin-wise average read counts were obtained from binary alignment map (BAM) files for each tumor and each matched normal sample (if available). Read counts were corrected for locus specific bias by subtracting from the vector of log-transformed reads its projection on the first *q* principal components found using log-transformed reads from a panel of normal samples.

### 2.3 Allelic copy ratio (BAF)

A SNP compendium was established using the dbSNP database^12^ and germline heterozygous SNPs were identified in each individual using `ModelSegments` in GATK. SNPs heterozygous in at least *k* = 5 matched normals were indexed in the following as *j* = 1,..., *M*. For each tumor sample the subset of heterozygous SNPs *j* were identified. Let 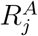 and 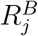 denote the corresponding allelic read depths, where A and B denote, respectively, the reference and alternative alleles according to dbSNP. Allelic read depths are assumed to follow Poisson distributions 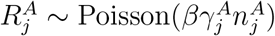 and 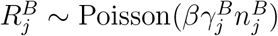,where *β* > 0 is the average read depth for the tumor sample, 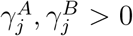 are allelic pref-erences and 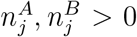 denote the number of copies of each variant allele in the tumor cells. Allelic preferences are platform dependent probabilities of a randomly chosen read fragment deriving from each of the two alleles, and we assume 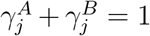. The B-allele frequency (BAF) of the *j*th SNP is 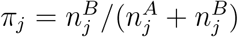 and is commonly estimated as

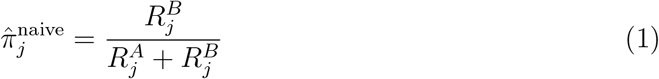

Conditional on the total read depth 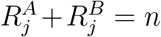, the distribution of 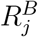 is binomial with parameters *n* and *p* = *b*/(*a* + *b*) where 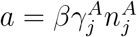 and 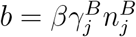. Thus, conditional on total read depth, we have 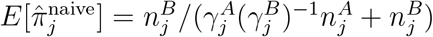 and the naive estimator is therefore conditionally unbiased only if 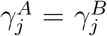. To avoid bias when there is an allelic preference 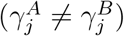, we propose the following adjusted estimate for π_*j*_:

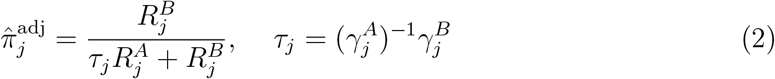

This estimator is easily verified to be conditionally unbiased for π_*j*_. Furthermore, the bias is negligible for moderate allelic preferences also in the unconditional case: if |*τ*_*j*_ − 1| *≤* θ, then to a second order approximation

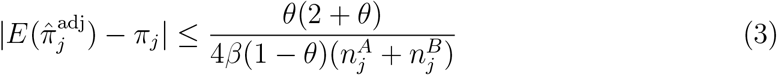

The bound in (3) is inversely proportional to the average read depth β and also to the total copy number. With no allelic preference, we have *τ*_*j*_ = 1, and we can set θ = 0 to obtain 0 on the right hand side of (3). With moderate allelic preference, we can set θ > 0 to be a small number, and the right hand side of (3) will be small. In order to compute the adjusted BAF estimate, we need to determine the allelic preferences. For this purpose, we use a panel of normal samples processed on the same sequencing platform. For a given SNP, suppose sequence data are available from *N* germline heterozygous normal samples. Suppressing the SNP index *j*, let 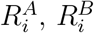 denote the allelic read depths and β_*i*_ the average read coverage for the *i*th normal sample (*i* = 1, 2,...,*N*). Since 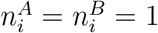 for all samples, we have

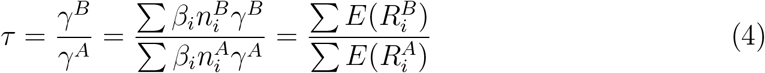

which immediately suggests the estimate 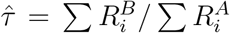. In summary, we propose the following BAF estimator that uses a panel of normal samples to adjust for allelic preference:

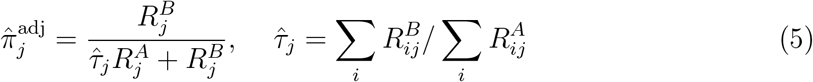

where 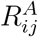 and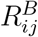 denote the allelic read depths for the *j*th SNP in the *i*th sample in the panel of normals.

### 2.4 Segmentation

Optimal segmentation of the genome requires use of both total copy ratio and BAF, since an allele-specific copy ratio event may affect both or only one of the two. A minor extension of the PCF algorithm^13^ that allows observations to be individually weighted was used for breakpoint detection. The weight of the *i*th point was chosen to be inversely proportional to the distance to the (*i* + 1)*th* data point. The algorithm is applied first to total copy ratio and subsequently to BAF to identify copy number neutral events (i.e. events that do not affect total copy number). For the latter, we use as input the estimates 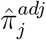 with allelic preferences chosen as in (4). Note that some elements of the sequence 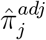reflect relative frequencies of the maternal allele and others the relative frequencies of the paternal allele (see section 2.5 for more details). For segments with allelic imbalance 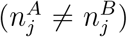 there will thus be two BAF bands symmetrically distributed around 0.5. To obtain a unimodal distribution, the BAF values are therefore first mapped to the values 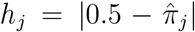. Sub-sampling is performed on the *h*_*j*_ values when the neighbouring distance is shorter than a threshold t_0_, to reduce serial correlation. In this paper *t*_0_ = 500 bp, approximately corresponding to the expected fragment length.

### 2.5 Phasing

The dbSNP-derived labels A and B have so far served the purpose of distinguishing two alleles of a biallelic SNP. These labels are unrelated to the physical location of the variants: some of the variants labeled A in a particular individual will be located on the maternal allele and some on the paternal allele. Hence, even within a particular copy segment, the B-allele frequencies 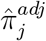 will be a mixture of maternal and paternal allele frequencies. For breakpoint detection (section 2.4) it suffices to collapse the two parental allele distributions to a common unimodal distribution by replacing 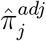 by its absolute deviation from 1/2. However, the mean of this unimodal distribution will deviate from the mean of either parental distribution when the latter overlap. For example, with two maternal alleles and one paternal allele, we would have (assuming for simplicity a normal distribution) 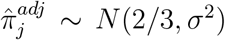 for some SNPs and 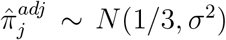 for others. For the unimodal distribution, we have 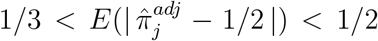 and with the mean increasing with the value of σ^2^. The two distributions would be increasingly difficult to identify with increasing allelic balance, and eventually the two bands would merge for allele-balanced data. Subtle allelic imbalances can occur due to subclonal copy number alteration or due to high infiltration of normal cells that dilute the 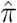 signal from the tumor cells.

Phasing aims to assign variants to their corresponding parental alleles, thus allowing more accurate estimation of parental allele frequencies^14,15^. Phasing leverages genetic linkage and haplotype information from large population databases to predict which heterozygous SNP variants are likely to occur on the same parental allele.

While phasing is challenging across entire chromosomes, it is feasible within shorter regions. In this study, we performed phasing using the SHAPEIT2 software^16^, based on inferred germline genotypes from matched normal samples. Haplotype data from the 1000 Genome Project phase 3 build 151^17^ was used for reference.

### 2.6 Determination of segment levels

For a locus *t*, we sought to estimate the total copy ratio *r*(*t*) and the minor parental allele frequency *q*(*t*). *r*(*t*) is estimated as the geometric mean of all the *R*_*i*_ values belonging to a given segment. Segment-wise allele frequencies are calculated as follows: Let 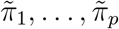 denote the phased estimated allele frequencies in a segment identified in section 2.3. Ideally, all the 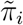 now correspond to one particular parental allele, and their average then reflect the parental allele frequency. In practice, the phasing algorithm will occasionally assign two neighbouring SNP variants to the same parental allele, whereas the correct solution would be to assign them to different ones, or vice versa. Such switch errors result in a switch of the parental assignment of all subsequent SNP variants. Consider a single segment identified by the method in section 2.3. If the phasing is correct, the parental allele frequency of the segment can be written as

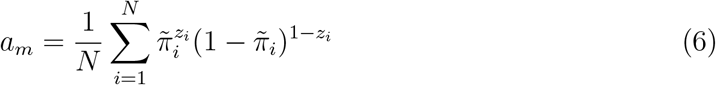

where *z*_*i*_ = 1 for all *i* and 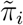 represents the relative frequency of the parental allele. However, if a switch error occurs at index *i*, the state of *z*_*i*_ needs to be flipped to the opposite state for all subsequent SNPs. This process must be performed iteratively for all switch errors in the segment (from left to right). In order to detect a switch error, we use the fact that the distribution of 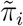 is different on each side of a switch error.More specifically, suppose the true allele frequency is π in the segment and that a switch error has occurred at the *i*th position. Then we have 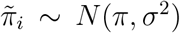 on the left side and 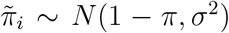 on the right side of the switch error. Hence, the magnitude of the expected shift in the mean of the distribution is |1 − 2π| and may be detected by segmentation. The PCF algorithm (see section 2.4) is used for this purpose. Occasionally, the iterative process described above reveals new switch errors not detected in the first pass. The process is therefore repeated a few times until convergence. In practice, one or two iterations will be sufficient to reach convergence. Note that a shift in the distribution will not be present if π = 1/2, but in this case a switch error will have no consequence for the estimated allele frequencies. For values of π very close to 1/2, the distribution shift may be too small to be detected by segmentation, but in this case the consequence for allele frequency estimation will be small. For any genomic locus s, the estimated minor parental allele frequency *q*(*s*) is defined as *min*(*a*, 1 − *a*) where a is the parental allele frequency of the segment that s belongs to.

Phasing can be computationally demanding and works best with a high genomic density of SNP variants. For whole-exome sequencing panels and other targeted panels, which generally have too low SNP densities for proper phasing, we use unphased allele frequencies to estimate parental allele frequencies. The first task is then to determine whether the allele frequencies from each segment represent a bimodal distribution (as expected for any unbalanced allele configuration), or a unimodal distribution (as expected for a balanced allele configuration). We expect a smaller proportion of the 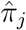 to be close to 0.5 in the former case than in the latter case. As the modes of a bimodal distribution will be mirrored around 0.5, the expression

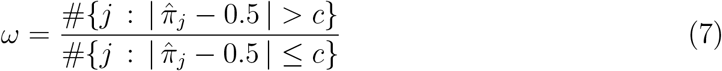

will take on larger values for a bimodal distribution of the 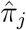 than for a unimodal distribution. The choice of c is somewhat arbitrary, but we would like a reasonable balance between the numerator and the denominator and use *c* = 0.5 *q*_0.90_ in this paper, where *q*_0.90_ is the 90th percentile of 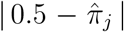. To summarize, we define the distribution of the 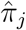 as bimodal if ω > ω^*^ for some threshold ω^*^ > 0. This threshold will depend on the number of data points 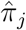 in a segment. We performed a simulation experiment to determine a proper threshold 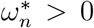 for each sample size *n* = 5, 6,..., *N* (sample sizes below five were considered too small to call bimodality). For each *n*, we performed simulations *b* = 1,…, *B* by first drawing independent samples 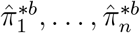 from a normal distribution with mean 0.5 and variance σ^2^ and then calculating ω^*\*b*^ using the formula in (7). Subsequently, the threshold 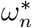 was defined as the 95th percentile of the numbers ω^*1^,..., ω^\**B*^. For a given segment, with *n* BAF probes, if 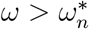 we call the distribution bimodal. Otherwise, we assume a unimodal distribution with mean 0.5 and estimate the standard error directly. Consider the case where the distribution is bimodal. We then model the observations 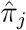 for a given segment as coming from a mixture of two normal distributions, where each reflects one of the two modes (the upper and lower branches of the BAF distribution, respectively). Assuming that the two modes have equal distance from 0.5, we thus have 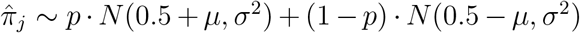 for some segment-specific *µ* and σ^2^. Note that the mixture coefficient *p* could take any value between 0 and 1, since the dbSNP-derived labels A and B are not random. We define the negative log-likelihood as

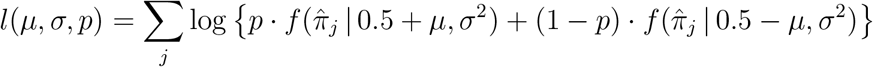

where *f*(*·*| *µ*, σ^2^) is the probability density of a normal distribution with mean *µ* and variance σ^2^. Maximizing *l*(*µ, σ, p*), we obtain parameter estimates 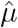, 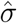, and 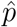. Only the first two of these values are used in practice; the latter is only a nuisance parameter.

### 2.7 Inference of discrete copy number states

Recall that *r*(*t*) and *q*(*s*) denote the estimated total copy ratio and minor parental allele frequency in locus *t* and *s*, respectively. The functions *r*(*t*) and *q*(*s*) generally reflect a mixture of tumor cells and normal cells and are related to the mean copy number *µ*(*t*) and the mean minor allele copy number *µ*^0^(*t*) in the tumor cell population by the equations (*s, t ∈ G*)

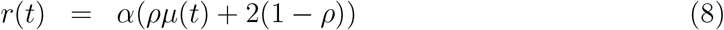

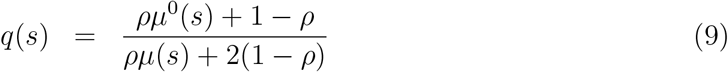

where ρ > 0 denotes the tumor cell proportion and α > 0 depends on the average sequencing coverage and the total DNA content of each tumor cell (i.e. the tumor cell ploidy). Both *ρ* and *α* are sample specific parameters and have to be determined from the data. Suppose *ρ* and *α* are determined through model selection (see below). Then Eqs. 8-9 can be used to derive *µ*(*t*), *µ*^0^(*t*) and *µ*^0^(*t*):

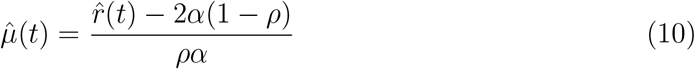

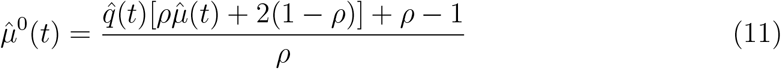

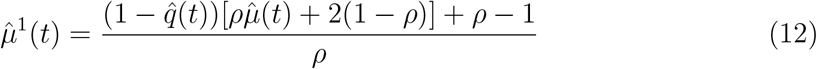

An approximate (1 −α) *·* 100% confidence interval for 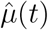 is 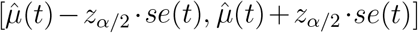 where *se*(*t*) is the standard error of 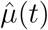. Likewise, an approximate (1 − α) *·* 100% confidence interval for 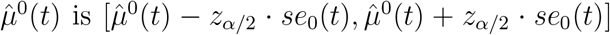 where *se*_0_(*t*) is the standard error of 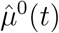. From Eq. 10 and Eq. we find that 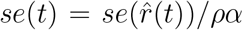.Furthermore, if we let 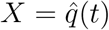 and 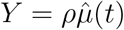, then:

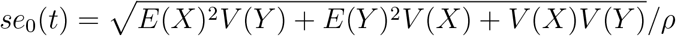

As the variance of 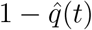 equals that of 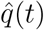, we may also assume that *se*_1_(*t*) = *se*_0_(*t*). As outlined above, *µ, µ*^0^ and *µ*^1^ may be estimated for any position *t* by Eq. 10-12, given the parameters *α* and *ρ*. Here we propose a selection criterion for *α* and *ρ*, which aims to achieve minor and major copy number estimates close to integer values. We minimize the loss function:

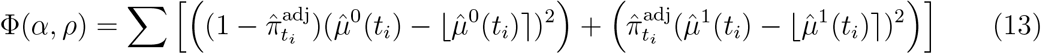

We determine optimal *α*and *ρ* values by calculating the above criterion for a grid of *α* and *ρ* values, defined within a biologically reasonable range. It can be shown that multiple solutions for *α* and *ρ* with very similar values for the criterion exist. E.g. for a truly diploid tumor, a solution for *α* and *ρ*corresponding to a doubling of the true ploidy will usually produce nearly equal values for the criterion. This is reflected in the regularity of local minima found across the Φ(*α, ρ*) plot. In order to select a reasonable model in situations with several local minima, we first determine all local minima over the defined range of combinations. We next compare the best solutions by a bootstrapping procedure, using the estimated distributions of 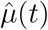, 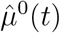 and 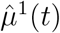 in each simulation, repeated for 100 iterations. For the final model selection, we choose the solution with the lowest ploidy, for which there are no significantly better solutions. A significant difference is determined if the bootstrap error distributions do not overlap at the 10th and 90th percentile for each of the two corresponding solutions. This implies that we always choose the simplest model supported by the data. Once tentative estimates for *α* and *ρ* are determined, we can estimate *µ, µ*^0^ and *µ*^1^ as well as their corresponding subclonality states. We define subclonal segments to encompass regions where the copy number estimates significantly deviate from integer values. The criteria for calling a segment subclonal are determined as a substantial deviation of the continuous copy number estimates 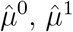 and 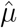 from an integer value (here defined as > 0.1), and a Bonferroni-adjusted p-value < 0.01.

### 2.8 Mutation multiplicity

A mutation initially present in one copy of DNA may be amplified by subsequent copy number alterations. The mutation multiplicity can in many cases be determined from allele-specific copy number estimates and mutation allele frequencies. To simplify the exposition, every position in the genome is assumed to mutate at most once, in line with the infinite sites model of molecular evolution^18^. Violations to this assumption are relatively rare; they can in some cases be detected, although we do not consider that here. The infinite sites model implies that at any given locus at most one of the two alleles is mutated, hence at a locus with *n*_*A*_ copies of allele A and *n*_*B*_ copies of allele B the mutation multiplicity is at most *max*(*n*_*A*_, *n*_*B*_).

Consider a biopsy with tumor cell fraction *ρ ∈* (0, 1] and suppose *n* > 0 somatic mutations are detected in copy number clonal genomic regions. A particular mutation will be carried by a proportion *α ∈* (0, 1] of the tumor cells, each containing c > 0 DNA copies of the locus, of which *k* > 0 copies harbor the mutation. All cells carrying the mutation are assumed to have the same number of mutant copies; hence *k* and c are uniquely defined for each mutation. The lemma below characterizes precisely what the possible mutation states (*α, k*) are for a genomic locus with a unique copy number state in the tumor cell population.

#### LEMMA 1

*Consider a mutation in a locus with total clonal copy number* c *>* 0 *in the tumor cell population; then the mutation state satisfies* (*α,k*) *∈ 𝒞where*

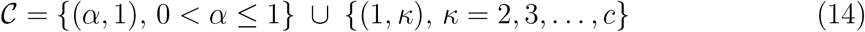

*Thus, the mutation is present in a subset of all tumor cells and in one single DNA copy per cell, or in all tumor cells and in one or more DNA copies per cell*.

PROOF

Let *C* denote the most recent common ancestor (MRCA) of the tumor cell population. Due to the copy number clonality assumption, a copy number alteration observed at present at this locus must have been present in *C*. If the mutation occurred in *C* or in one of its descendants, only one copy of DNA can be affected (otherwise, the mutation must have occurred multiple times). Hence, we have (*α, k*) = (*α*, 1) for some *α ∈* (0, 1]. If, on the other hand, the mutation occurred in an ascendant of C, then the mutation is present in all tumor cells and consequently we have (*α, k*) = (1, *k*) for some *k∈ {*1, 2, 3,…, *c}*. Hence, we must have (*α, k*) *∈ 𝒞*.

Combined, the mutation state (*α, k*) and the total copy number *c* uniquely determine the variant allele fraction of a mutation. To see this, consider a population of *N* cells. The number of aberrant cells is *ρN*, the number of cells carrying the mutation is *αρN*, and the total number of mutant DNA copies is *αkρN*. On the other hand, the total number of DNA copies is *cρN* + 2(1 *− ρ*)*N*, so the proportion of mutant DNA copies (i.e. the variant allele fraction) is

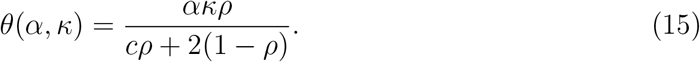

In practice, the values of *α* and *k* are unknown. However, we can determine the proportion of mutant DNA copies *θ* (*α, k*) from a sequencing experiment, and the aberrant cell fraction *ρ* and the total copy number *c* can be determined from the previously described copy number analysis. When used in combination, this allows us to determine the value of the product *αk* from eq. (15). More importantly, the next lemma shows that *α* and *k* are both uniquely determined from (15).

#### LEMMA 2

*Let ρ* > 0 *denote the aberrant cell fraction and consider a mutation at a locus with clonal total copy number c* > 0 *in the tumor. Suppose* θ : *𝒞*→ **R** *is defined as in (15) and let*

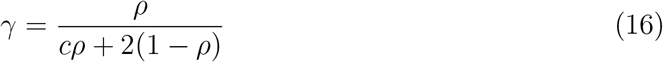

*Then* θ *is a bijection between 𝒞 and 𝒟* = (0, γ] *⋃ {*2γ, 3γ, …, cγ*}. In particular, for any*

*z ∈ 𝒟 there is a unique* (*α, k*) *∈ 𝒞 such that* θ (*α, k*) = *z*.

PROOF

If (*α, k*) *∈ 𝒞* then *θ* (*α, k*) = *αkγ*. Since *α, k* > 0 and also *γ* > 0, we must have θ (*α, k*) > 0. Suppose *θ* (α ′, *k*′)) for (*α, k*), (α ′, *k*′) ∈ *𝒞*. Then *αk* γ= α ′, *k*′ γ, which implies that *αk*= α ′, *k*′, since γ *≠*0. If *α, k* ≤ 1, then according to Lemma 1 we must have *k* = 1 and also *k*′ = 1, but then we must also have *α*= α ′. If, on the other hand *α k* > 1, then according to Lemma 1 we must have *α* = α′= 1 and it follows that *k* = *k*′. In both cases, we have (*α, k*) = (α ′, *k*′), hence *θ* is one-to-one. Furthermore, if *α k* ≤ 1, we have θ (*α, k*) = *α k* γ ≤ γ, and if *α k* > 1 then θ (*α, k*) = γ ≤ γ =≤ γ, where (according to Lemma 1) *k* = 1, 2,..., c. Hence, the image of 𝒞 under the map θ is 𝒟. This concludes the proof.

We now make use of the two lemmas above to derive an efficient algorithm for determination of the mutation state (α, *k*) for a set of mutations. Suppose we have n mutations with variant read depths *r*_1_, …, *r*_*n*_ and total read depths *R*_1_, …, *R*_*n*_. Conditional on the total read depths, we model the variant read depths as binomial variables,

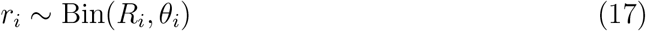

where θ_*i*_ > 0 is the proportion of the DNA sequences in the sample that harbors the ith mutation. This proportion is determined by the mutation state (α_*i*_, *k*_*i*_) for the ith mutation and the total copy number c_*i*_ through equation (15), i.e. we have θ_*i*_ = θ (α_*i*_, *k*_*i*_; *c*_*i*_) where the third argument emphasizes that the constant c in (15) is equal to c_*i*_. Let **X** denote the observed data (*r*_*i*_, *R*_*i*_) for all mutations *i* = 1, …, *n* and let *k* = (*k*_1_, …, *k*_*n*_) and *k* = (*k*_1_, …, *k*_*n*_). Then the likelihood becomes

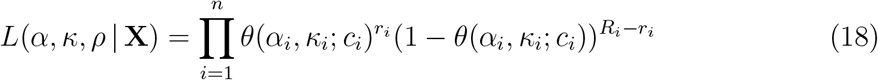

where ρ denotes aberrant cell fraction as before. We first consider the problem of maximizing the likelihood (18) with respect to *α* and *k* for fixed value of ρ. This can be achieved by maximizing separately each term in the product, or equivalently maximizing the log-transformed and scaled terms

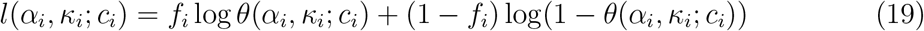

over (*α*_*i*_, *k*_*i*_) ∈ *𝒞*_*i*_, where *f*_*i*_ = *r*_*i*_/*R*_*i*_. The function *h*(θ) = *f* log *θ*+(1*−f*) log(1−*θ*) is easily seen to satisfy h″ (*f*) = 0 and h″ (θ) < 0. Hence, *h*(θ) is strictly concave with a unique global maximum in θ = *f*. Thus, if there exists (*α*_*i*_, *k*_*i*_) *∈ 𝒞*_*i*_ such that θ(*α*_*i*_, *k*_*i*_; *c*_*i*_) = *f*_*i*_, then this is the maximizer of (19). Recall from Lemma 2 that the image of θ(*·, ·*; *c*_*i*_) is *𝒟*_*i*_ = (0, *γ*_*i*_] ⋃ *{*2*γ*_*i*_, 3*γ*_*i*_,..., c_*i*_*γ*_*i*_*}*. If *f*_*i*_ *≤ γ*_*i*_ then according to the proof of Lemma 2 we must have *α*_*i*_ = *f*_*i*_ and *k*_*i*_ = 1. If *f*_*i*_ ∈ (*kγ*_*i*_, (*k* + 1)*γ*_*i*_], for some *k* = 1, 2,..., *c*_*i*_ − 1, the maximum is obtained at either *kγ*_*i*_ or (*k* + 1)*γ*_*i*_. The solution is then either (again according to the proof of Lemma 2) *α*_*i*_ = 1, *k*_*i*_ = *k* or *α*_*i*_ = 1, *k*_*i*_ = *k* + 1. To determine which of the two candidates forms the solution, we compute (19) for both and select the one with highest value. If *f*_*i*_ > *c*_*i*_ *γ*_*i*_, then the maximum is obtained at *c*_*i*_*γ*_*i*_ and the solution is *α*_*i*_ = 1 and *k*_*i*_ = *c*_*i*_. The above discussion suggests the following o(n) algorithm to optimize the likelihood in (18).

#### Algorithm 1: Mutation counting

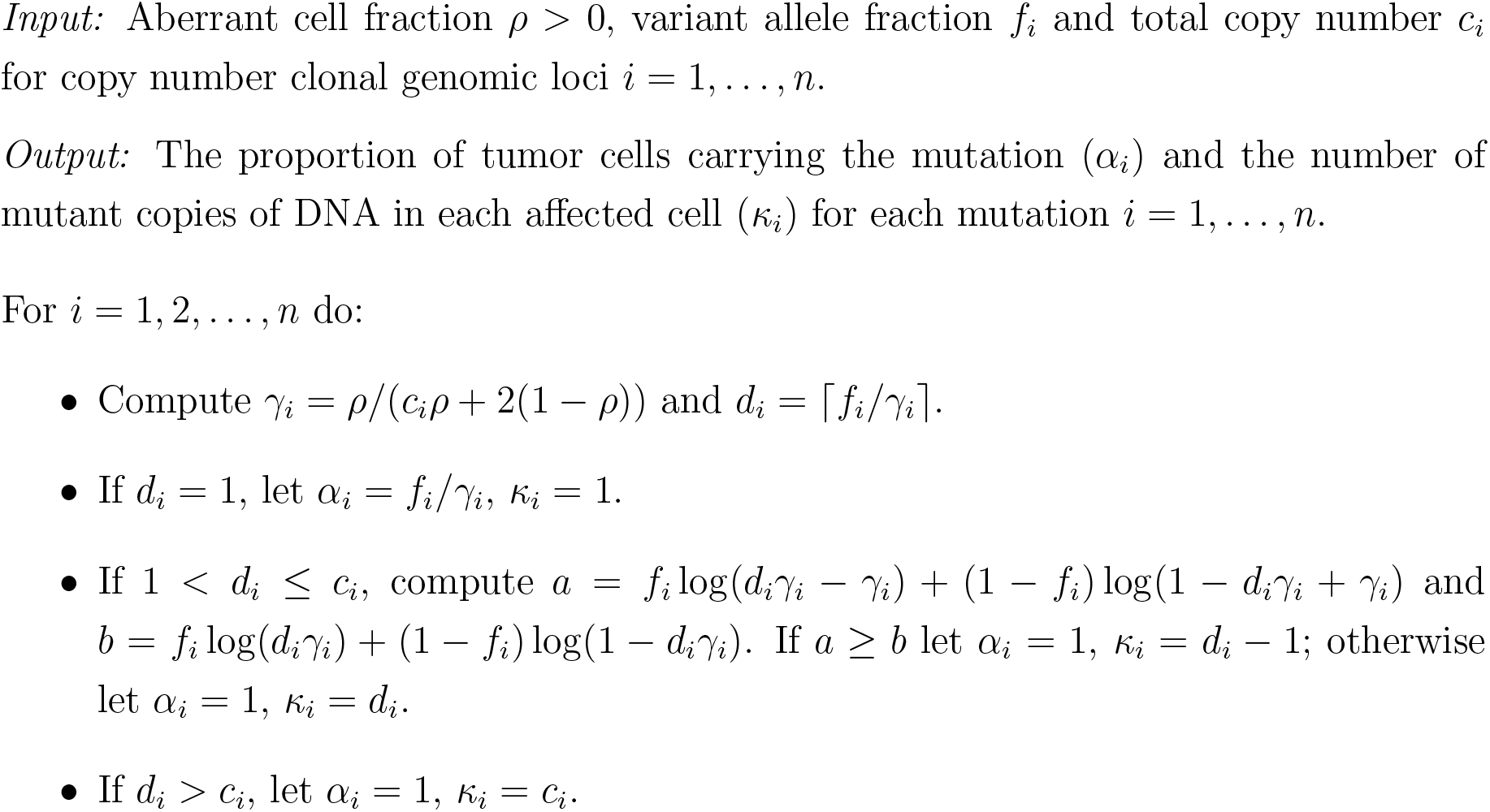

### 2.9 Mutation counting with unknown tumor percentage

The algorithm described in the previous section assumes that the aberrant cell fractionρ is given. If is ρunknown, we may maximize the likelihood in Eq. over (*α, k*, ρ) by applying Algorithm 1 repeatedly with ρ selected from a grid of possible values 0 *<ρ*_1_ *<*ρ_2_ *< … <* ρ_*s*_ ≤1. We then select the grid value *ρ*_*j*_ that maximizes the likelihood with respect to the remaining parameters *α* and *k*.

### 2.10 Mutation counting under subclonal copy number

The model presented above and Algorithm 1 rely on the assumption of mutations being located in copy number clonal regions. In practice, this entails filtering out mutations residing in copy number subclonal regions prior to the application of Algorithm 1. To investigate the effect of subclonal copy number on the mutation counting, suppose the *i*th locus exists in three different copy number states: 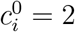 (corresponding to the infiltrating normal cells), 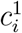 and 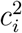. In the following, we refer to the cells with copy number 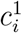 as subclone 1 and the cells with copy number 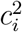 as subclone 2. Let 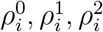 denote the corresponding proportions of cells in the sample, such that 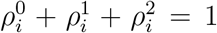. Let 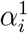 and 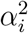 be the proportion of cells carrying the *i*th mutation in the two subclones and let

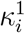and 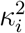 denote the number of mutant copies of DNA in each affected cell in the two subclones. By the same reasoning as before, it is readily shown that 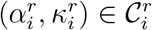 where

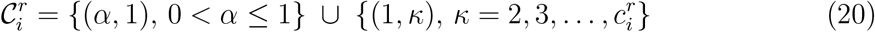

where r = 1, 2. In a sample of N cells, the number of mutant DNA copies is now 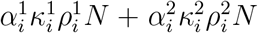and there are 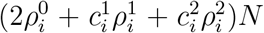N copies of DNA in total. The proportion of mutant DNA at the *i*th locus is thus

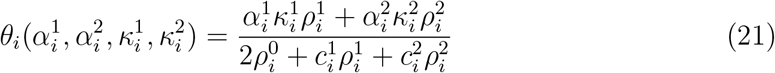

In contrast to the monoclonal case described by Eq. 15, the subclonal case described by Eq. 21 is not injective. Hence, in general there is not a unique set of parameter values for 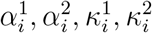that maximizes the corresponding likelihood analogue of Eq. 18.

### 2.11 Material and sequence analysis

#### The Cancer Genome Atlas Program (TCGA)

DNA sequencing data were obtained from a subset of 100 breast cancer patients from the TCGA cohort^19^. Informed consent was obtained from participants; data access and use followed TCGA policies. DNA was extracted from fresh frozen tumor tissue and matched normal samples. DNA whole exome sequencing data was available from all 100 patients; for five patients, whole genome sequencing data from the corresponding tumor samples and matched normal samples were also available. Six samples were excluded from further analyses due to evidence of low tumor purity based on copy number analysis.

#### IMPRESS-Norway

DNA was extracted from fresh frozen tumor tissue and matched normal tissue (blood) for 23 patients in the IMPRESS-Norway study^20^ (Improving public cancer care by implementing precision medicine in Norway, NCT04817956). The study has been approved by the Regional Committee for Medical and Health Research Ethics for southeast Norway (approval number 200764). Patients received written and oral information about the study and signed informed consent. The QIAamp DNA Micro Kit (Cat. No. 56304) was used for isolation of DNA from fresh frozen tissue; the EZ2 350 µl Blood kit (Cat. No 951054) was used for isolation of DNA from EDTA blood samples. DNA purity was assessed using the NanoDrop One machine; the concentration was determined using the Qubit 4 Fluorometer. DNA from FFPE tissue from six separate samples was extracted using the AllPrep DNA/RNA FFPE Kit (50) (Cat. No 80234). WGS DNA libraries were prepared using the Illumina DNA PCR-Free Prep Tagmentation kit and sequenced on the NovaSeq 6000 machine; alignment to the GRCh38 reference genome was performed using BWA-MEM. TSO500 DNA libraries were prepared using the Illumina TruSight Oncology 500 High-Througput kit and sequenced on Illumina NextSeq 500 and NovaSeq 6000 machines; alignment to the GRCh37 reference genome was performed using BWA-MEM.

### 2.12 Copy number analysis

WGS-based ASCAT results from the TCGA dataset were obtained from previously published supplementary data^21^. WGS-based ASCAT results from the IMPRESS-Norway study were generated using the Sarek pipeline (version 3.1.2)^22^ with ASCAT (version 3.0.0)^23^, applied to paired tumor-normal BAM files under default settings. WES-based ASCAT results for the TCGA dataset were obtained using the Sarek pipeline (version 3.1.1) with ASCAT (version 3.0.0)^23^, applied to paired tumor-normal BAM files under default settings. For ASCAT analyses of WES data, reference files for GC correction and replication timing were prepared following the recommendations in the Sarek documentation (https://nf-co.re/sarek/3.4.0/docs/usage).

Copy number results from PURPLE (version 4.0)^8^, based on WGS data, were generated using the `hmftools` pipeline (https://github.com/hartwigmedical/hmftools) with default settings. Allele frequencies and read depth ratios were obtained from AMBER (version 3.9.1) and COBALT (version 1.15.2), respectively^24^.

FACETS^9^ results based on WES data were obtained using the R package `facetsSuite` (version 2.0.8), along with wrapper scripts provided on the `facetsSuite` GitHub repository (https://github.com/mskcc/facets-suite). Allele frequencies were calculated using the `snp-pileup` function, which is implemented within the `facetsSuite` package.

PureCN analyses of TSO500 data were performed using PureCN (version 2.8.1)^7^, executed via a Docker container (tag: purecn:2.8.1-amd64). PureCN was run with the option `--fun-segmentation none` on segmentation data generated with GATK (version 4.2.0.0) from LocalApp (version 2.2.0.12) output.

Total copy ratios and allele frequencies obtained using GATK (version 4.2.0.0) served as input data for the CANCAN algorithm. For the WGS data, read counts were collected across static 1 kb windows. For the WES and TSO500 data, platform-specific targets were used to define the genomic windows from which read counts were collected. Additionally, in the TSO500 data, genomic windows were extended up to 250 bp on each side of the platform-specific targets (see Supplementary for details). The `CollectReadCounts` GATK tool was used to extract read counts, and normalization was performed with the `DenoiseReadCounts` tool, which relies on a panel of normal samples (PoN). For WGS data from the IMPRESS-Norway study, a PoN was constructed using 82 normal samples, whereas WGS data from the TCGA study were normalized using a single matched normal sample. In the WES data, the PoN was built from 100 matched normal samples. For the TSO500 data, 102 normal samples from the IMPRESS-Norway study were used to build a PoN. Germline heterozygous SNPs were identified in each individual using GATK’s `ModelSegments` tool. Only data from chromosomes 1-22 were included in the analysis. The segmentation parameter lambda was set to 100 for WGS and WES data, and to 15 for the TSO500 data. The minimum tumor purity was set to 15%. GRCh37-based genomic positions were converted to the GRCh38 reference genome for the purpose of integrated analyses, using the `segment liftover` software (https://github.com/baudisgroup/segment-liftover).

### 2.13 Estimation of DNA ploidy using image cytometry

Samples were prepared from FFPE specimens; monolayers of cell nuclei were prepared from one or more 50-µm sections cut from the blocks and stained with Feulgen–Schiff according to established protocols^25,26^. Automated image-based DNA cytometry was performed using Ploidy Work Station (PWS, Room 4 Ltd.) together with a Zeiss Axio-Plan light microscope with a 546-nm green filter and a black and white digital camera (AxioCam MRm, Zeiss). Automated imaging was followed by automatic segmentation of cell nuclei in the captured microscope images. Segmented cell nuclei were classified using PWS and trained personnel classified DNA ploidy results as diploid or aneuploid.

## 3 Results

### 3.1 CANCAN output

The main output of CANCAN is an estimate of the allele-specific absolute copy number, along with corresponding estimates of purity and ploidy (Figure 1). CANCAN assesses subclonality within each segment and reports p-values for the allele-specific copy number estimates. To support interpretation, CANCAN visualizes the final likelihood in a sunrise plot enabling direct comparison of likelihoods across alternative purity–ploidy solutions. It further stratifies results by minor and major alleles, facilitating evaluation of each allele’s contribution to the overall likelihood. CANCAN generates a complete results plot for a selected number of best-scoring solutions. Additionally, CANCAN provides optional and detailed visualizations for variants and genes of interest. These plots enable in-depth assessment of the local genomic context for biomarkers of interest (Figure 2).

**Figure 1.**
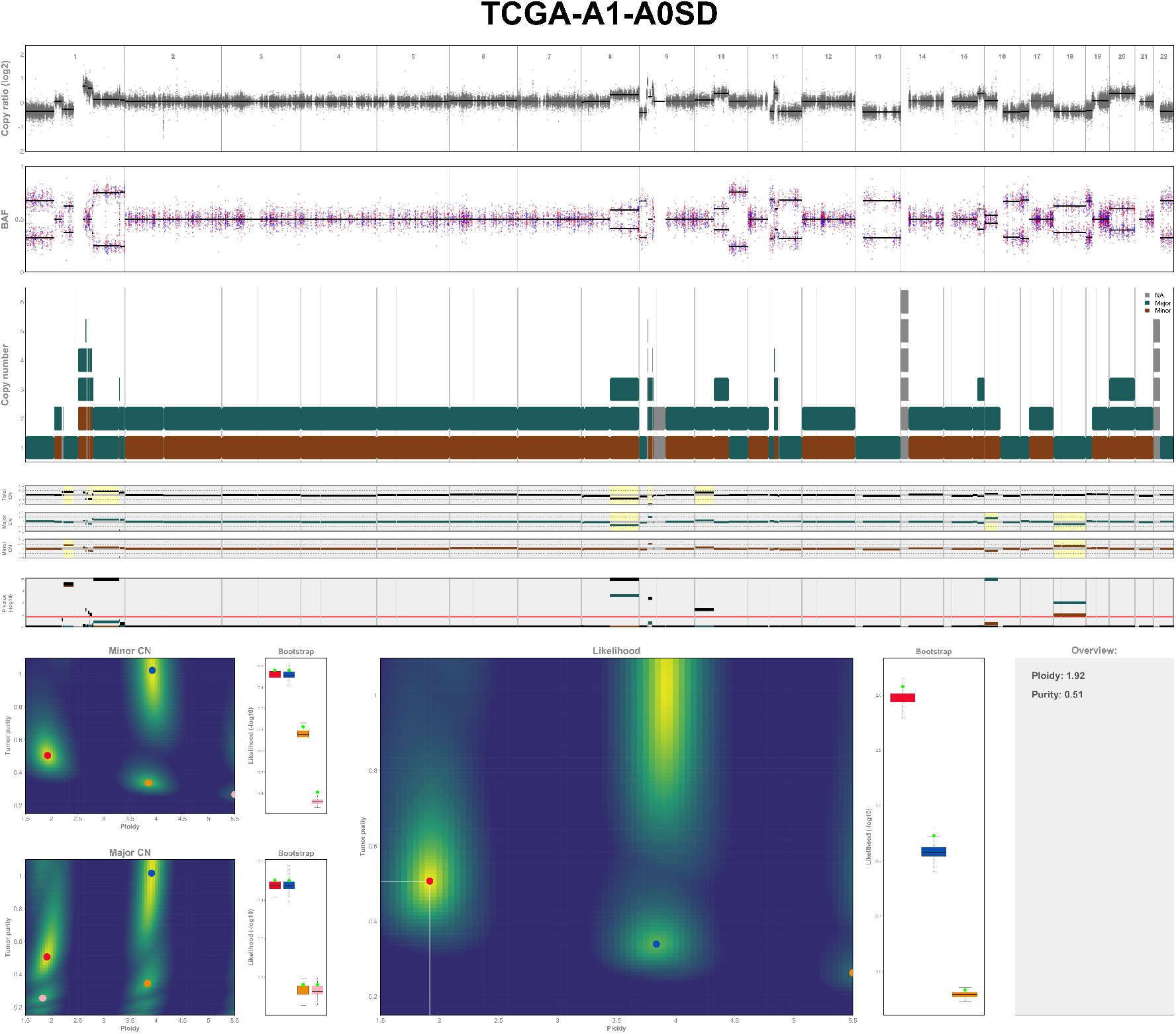
Main result plot generated by CANCAN. The first two tracks show the normalized input copy-ratio data and B-allele frequency (BAF), respectively. Both plots show the entire genome, ranging from chromosome one on the left side, to chromosome 22 on the right side. The horizontal black lines are the resulting segmentation from CANCAN. The blue and red dots in the BAF-plot represent the reference and alternative allele frequencies, respecitvely, of each heterozygous SNP in the data. The third track displays the allele-specific copy-number profile generated by CANCAN, where the green color represents the major allele and the brown color represents the minor allele. Grey indicates that the algorithm could not determine the minor or major component, only the absolute copy number. The subsequent track presents the non-rounded copy-number estimates for each allele, with segments classified as subclonal highlighted in yellow. The final track shows the corresponding p-values. The three sunrise plots illustrating the likelihood landscape across alternative purity–ploidy solutions are shown at the bottom, with contributions from the minor CN and major CN alleles displayed separately. A direct comparison of alternative purity–ploidy solutions derived from bootstrapping, repeated for 100 iterations, is shown on the right, where the red box-plot is the solution with highest likelihood. Lastly, in the overview, the final model is presented with its estimated ploidy and purity. For the final model selection, we choose the solution with the lowest ploidy, for which there are no significantly better solutions, as explained i2n2the method section.

**Figure 2.**
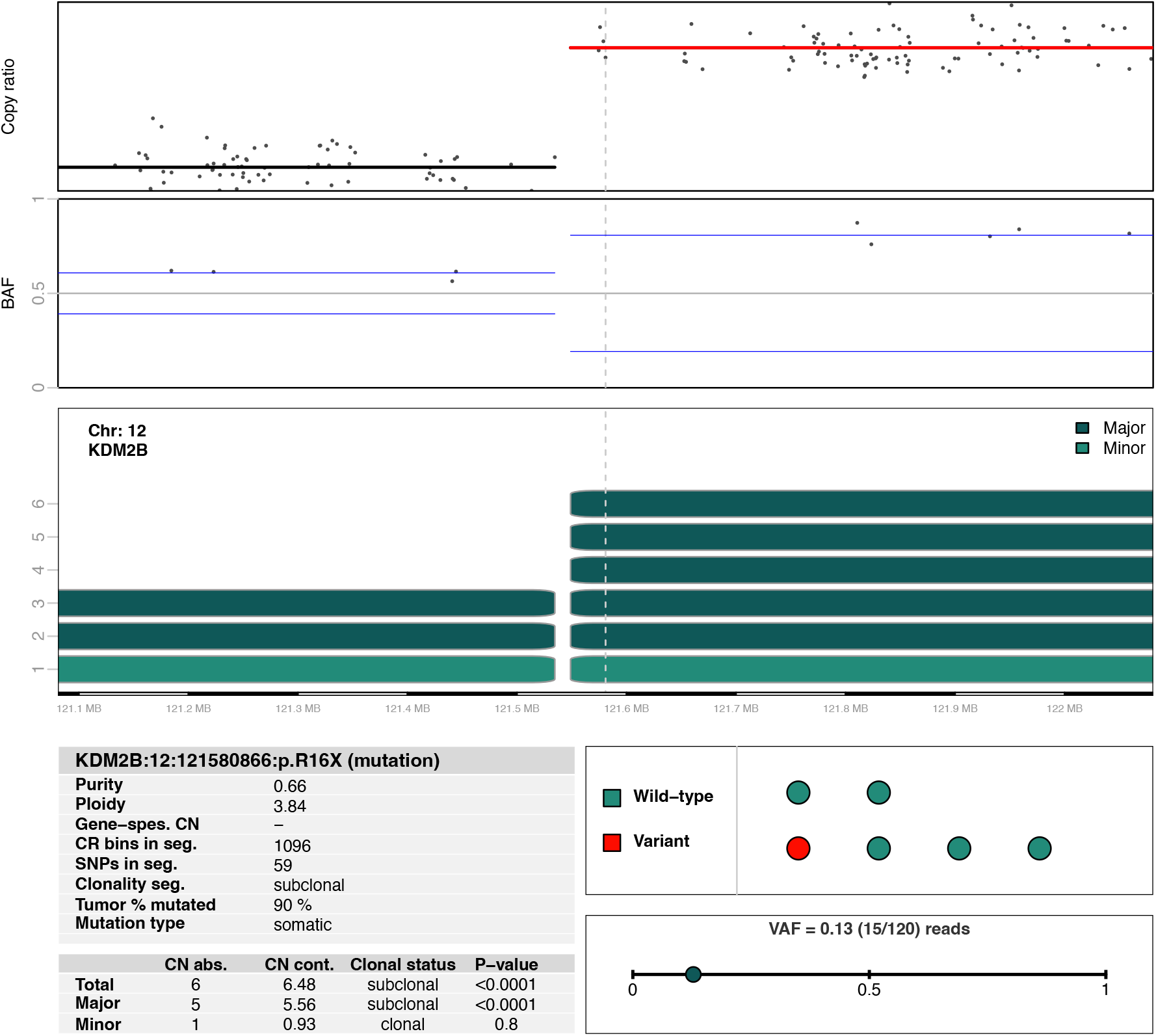
Example of a detailed visualization of the genomic context of a mutation of interest. The plot allows in-depth assessment of the local genomic context surrounding the biomarker of interest (position depicted by the dashed vertical line). The first two tracks show normalized input copy-ratio data and B-allele frequency (BAF) within a narrow genomic window, with estimated averages indicated by segment lines. The third track displays the corresponding allele-specific copy-number estimates. Subsequent windows present estimates of subclonality and the number of alleles affected by the variant.

### 3.2 Benchmarking purity and ploidy estimates

CANCAN’s performance was benchmarked against other high-performance copy number methods, utilizing sequencing data from 94 TCGA samples and 23 IMPRESS-Norway study samples. Whole-genome sequencing (WGS) data were available for 28 samples, whole-exome sequencing (WES) for 94 samples, and targeted sequencing (TSO500) for 23 samples (Table 1). Each method was applied only to the sequencing platforms for which it was primarily designed. For WGS data, we used CANCAN, ASCAT^6,27^, and PURPLE^8^; for WES data we used CANCAN, ASCAT, and FACETS^9^; and for TSO500 data we used CANCAN and PureCN^7^ (Table 2). Two central parameters in copy number estimation are tumor purity and tumor ploidy. We initially focused on comparing these parameters across methods. For this purpose, we considered two methods to be concordant when their ploidy estimates differed by less than 0.5 and their purity estimates differed by less than 0.1. Across the WGS cohort, CANCAN demonstrated concordance with ASCAT and PURPLE in all samples where ASCAT and PURPLE produced concordant results (17/28) (Table 3). In the 11 samples where ASCAT and PURPLE were discordant, CANCAN’s estimates aligned with one of the two methods in every case. In the WES cohort, CANCAN showed concordance with ASCAT and FACETS in 51 of the 55 samples (93%) where ASCAT and FACETS were concordant (55/94). As previously explained, CANCAN provides ranked alternative solutions for purity and ploidy. When the best alternative solution from CANCAN was included, the concordance increased to 98% (54/55). Among the samples where ASCAT and FACETS produced discordant results (39/94), CANCAN’s estimates were consistent with one of the two methods in 82% (32/39) of the cases, increasing to 92% (36/39) when the best alternative solution from CANCAN was included.

**Table 1.** Overview of the number of patients included in this study, stratified by sequencing platform and study.

|  | **All** | **TCGA** | **IMPRESS** |
| --- | --- | --- | --- |
| **WGS** | 28 | 5 | 23 |
| **WES** | 94 | 94 | 0 |
| **TSO500** | 23 | 0 | 23 |

**Table 2.** Overview of the copy number methods applied to each sequencing platform.

|  | **CANCAN** | **ASCAT** | **PURPLE** | **FACETS** | **PureCN** |
| --- | --- | --- | --- | --- | --- |
| **WGS** | **X** | **X** | **X** |  |  |
| **WES** | **X** | **X** |  | **X** |  |
| **TSO500** | **X** |  |  |  | **X** |

**Table 3.** Overview of consensus rates for purity and ploidy estimates in the WGS and WES datasets.

|  |  |
| --- | --- |
| **WGS - ploidy & purity** | **n (%)** |
| **ASCAT & PURPLE agreement** | **17/28 (61)** |
| CANCAN agreement | 17/17 (100) |
| **ASCAT & PURPLE disagreement** | **11/28 (39)** |
| CANCAN & ASCAT agreement | 7/11 (64) |
| CANCAN & PURPLE agreement | 5/11 (45) |
| CANCAN & ASCAT or CANCAN & PURPLE agreement | 11/11 (100) |
| **WES - ploidy & purity** | **n (%)** |
| **ASCAT & FACETS agreement** | **55/94 (59)** |
| CANCAN agreement | 51/55 (93) |
| **ASCAT & FACETS disagreement** | **39/94 (41)** |
| CANCAN & ASCAT agreement | 28/39 (72) |
| CANCAN & FACETS agreement | 7/39 (18) |
| CANCAN & ASCAT or CANCAN & FACETS agreement | 32/39 (82) |

For all samples with TSO500 data, corresponding WGS data were available. Bench-marking of copy number estimates from TSO500 data was performed by comparing ploidy and purity estimates derived from TSO500 with those obtained from WGS, which served as the gold standard. To avoid self-reference, WGS-based estimates were derived using ASCAT and PURPLE, and only the 16 samples in which these two methods were concordant were included in the analysis.

In all of these cases, TSO500-based CANCAN estimates were consistent with matched WGS results, with median differences in purity and ploidy of 2% and 0.03, respectively (Table 4, Supplementary Figure 1). In comparison, TSO500-based PureCN estimates were consistent in 81% (13/16) of cases. The mean coverage of the original TSO500 sequencing data was 736x. To assess robustness under reduced coverage, the sequencing data were randomly down-sampled *in silico* to average coverages of 300x and 100x. At 300x coverage, CANCAN achieved 81% (13/16) concordance with WGS estimates, and at 100x coverage, 62% (10/16) concordance was achieved. For PureCN, concordance was 81% (13/16) at 300x and 69% (11/16) at 100x (Table 4).

**Table 4.** Overview of consensus rates for purity and ploidy estimates in the TSO500 dataset.

|  | **Full coverage** | **300 X** | **100 X** |
| --- | --- | --- | --- |
| CANCAN TSO500 & WGS | 16/16 (100) | 13/16 (81) | 10/16 (62) |
| PureCN TSO500 & WGS | 13/16 (81) | 13/16 (81) | 11/16 (69) |

### 3.3 Benchmarking copy number estimates

To benchmark allele-specific absolute copy number estimates, a metric was defined to quantify differences between methods. The metric was defined as the average absolute difference in copy number between two methods, where 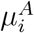 and 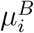 define the total copy number estimates for methods *A* and *B* at a genomic position *i*, sampled at evenly spaced genomic positions:

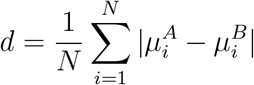

In this study *N* = 30309, corresponding to positions spaced every 10 kilobases across the genome. When comparing copy number estimates derived from WGS data, the median difference in copy number, as measured by d, was 0.10 for CANCAN relative to ASCAT, and 0.03 relative to PURPLE(Table 5, Figure 3, Supplementary Figure 2). In contrast, the median difference between ASCAT and PURPLE was 0.17. For WES data, the median d for CANCAN was 0.10 relative to ASCAT and 0.47 relative to FACETS, while the median difference between ASCAT and FACETS was 0.51 (Table 5, Figure 3, Supplementary Figure 3). For the TSO500 dataset, d values were calculated by comparing TSO500-based copy number estimates with matched WGS-based estimates, considering only regions with concordant copy number calls across ASCAT and PURPLE. The median difference between WGS- and TSO500-based estimates measured by d was 0.03 for CANCAN and 0.06 for PureCN.

**Table 5.** Median differences in copy number estimates determined by the metric *d*.

| **WGS** | **CANCAN vs ASCAT** | **CANCAN vs PURPLE** | **ASCAT vs PURPLE** |
| --- | --- | --- | --- |
| Difference (d) | 0.10 | 0.03 | 0.17 |

| **WES** | **CANCAN vs ASCAT** | **CANCAN vs FACETS** | **ASCAT vs FACETS** |
| --- | --- | --- | --- |
| Difference (d) | 0.10 | 0.47 | 0.51 |

| **TSO500** | **CANCAN vs WGS** | **PureCN vs WGS** |
| --- | --- | --- |
| Difference (d) | 0.03 | 0.06 |

**Figure 3.**
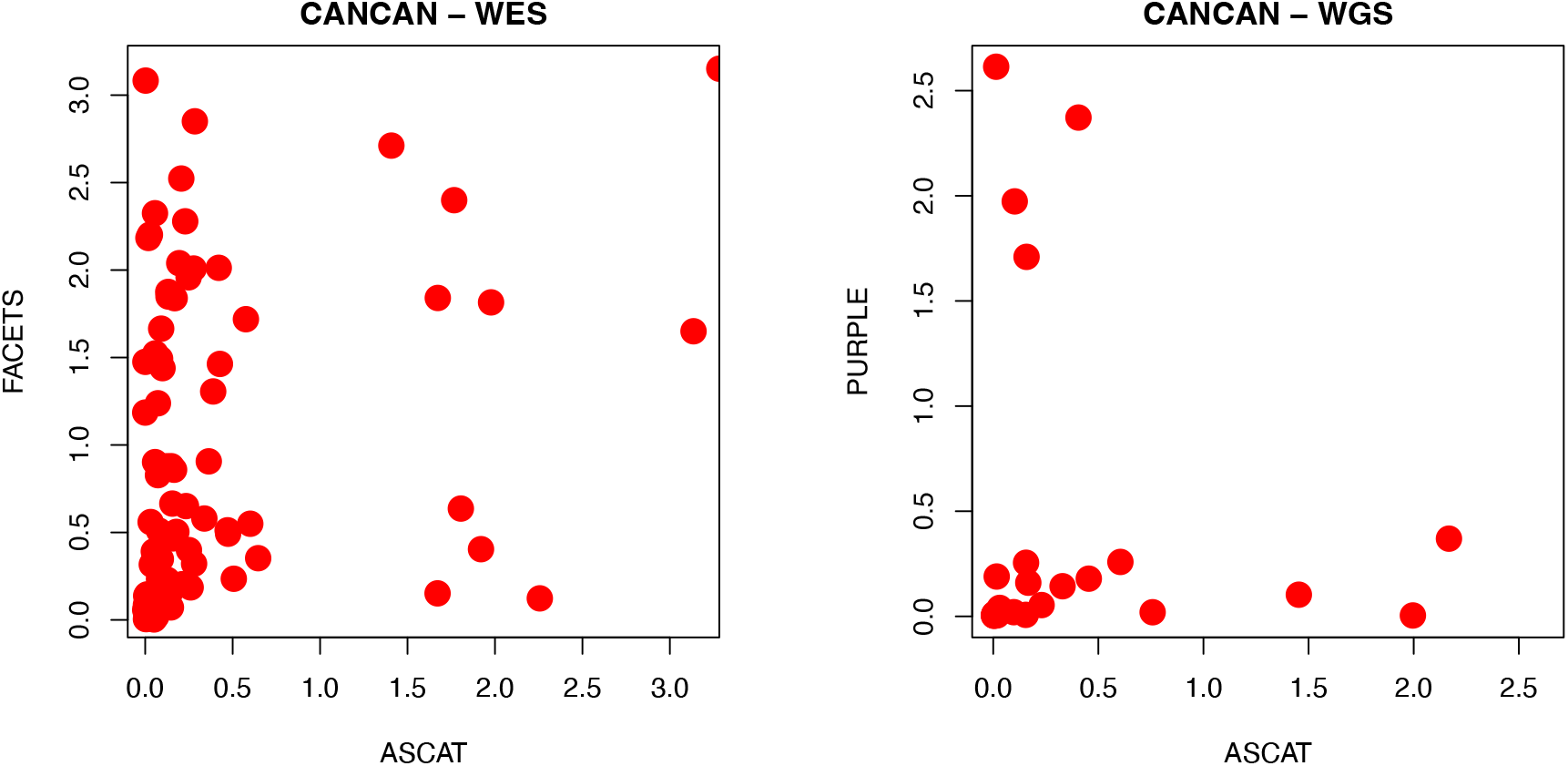
Differences in WES and WGS-based copy number estimates according to the metric *d*, comparing results from CANCAN with alternative best-practice methods.

### 3.4 Integrated optical density analysis

TSO500 data from additional FFPE samples were available for six patients included in the IMPRESS-Norway study, and copy number profiles were generated using CANCAN for these samples. Independent ploidy estimates were obtained via integrated optical density (IOD) analysis, and concordance with CANCAN-based estimates was evaluated (Figure 4, Table 6). One of the six samples showed low tumor purity and unreliable copy number results and was excluded from further analysis. Among the remaining five samples, three showed nearly identical ploidy estimates between CANCAN and IOD. The fourth sample (IPD0288) displayed concordant results when considering CANCAN’s second-ranked ploidy estimate. The fifth sample (IPD0661) did not show concordant ploidy estimates for either the top-ranked or second-ranked CANCAN solution. Purity estimates based on CANCAN were relatively low (24%) for this sample, and WGS-based CANCAN results from a separate biopsy indicated extensive subclonality. Ploidy estimates from PureCN were generally consistent with those from CANCAN; however, for sample IPD0288, neither the top-ranked nor second-ranked PureCN solution matched the IOD estimate.

**Table 6.** Overview of ploidy estimate correspondence, comparing IOD-based estimates with ploidy estimates from CANCAN and PureCN, respectively, showing ploidy estimates for the two best solutions. CANCAN and PureCN were applied to TSO500 sequencing data from FFPE tissue.

| **Sample ID** | **IOD** | **CANCAN**<br>**1st solution** | **CANCAN**<br>**2nd solution** | **PureCN**<br>**1st solution** |
| --- | --- | --- | --- | --- |
| IPD0072 | 1,60 | 1,56 | 5,15 | 1,54 |
| IPD0612 | 1,82 | 1,78 | 3,66 | 1,81 |
| IPD0823 | 3,90 | 3,85 | 1,87 | 3,94 |
| IPD0288 | 2,62 | 5,26 | 2,65 | 4,86 |
| IPD0661 | 3,12 | 1,78 | 3,78 | 2,56 |

**Figure 4.**
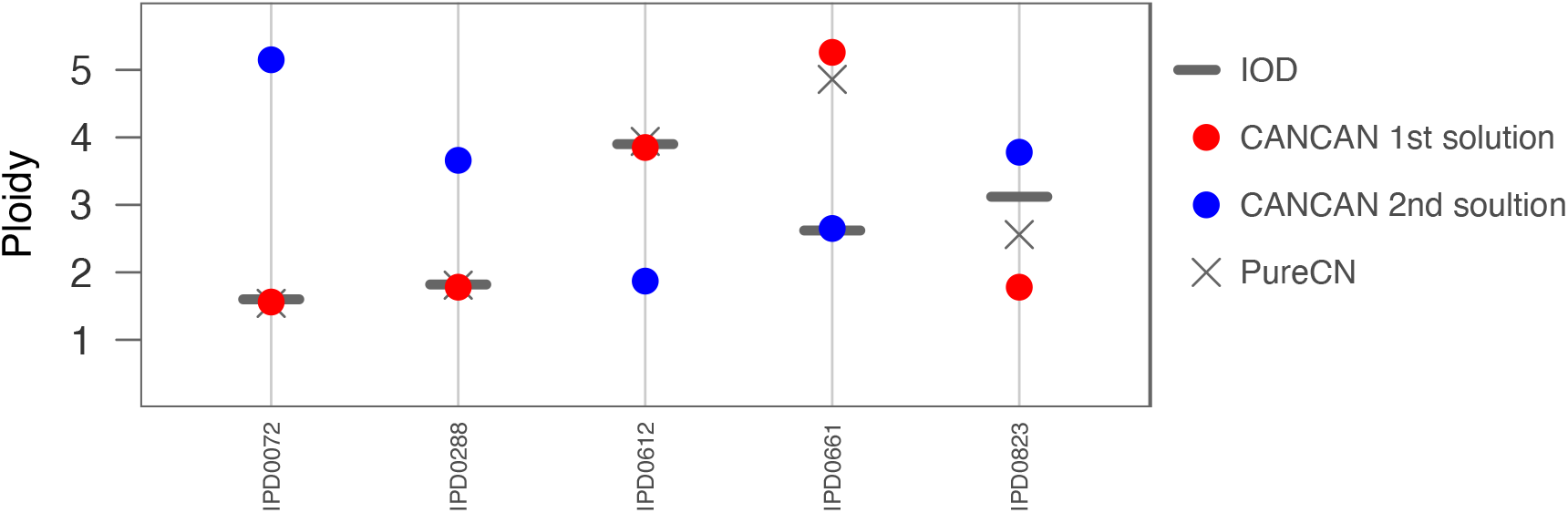
Comparison of ploidy estimates obtained from integrated optical density (IOD) to ploidy estimates generated by CANCAN and PureCN based on TSO500 sequencing of the matched FFPE tissue.

## 4 Discussion

A key strength of CANCAN is its ability to analyze sparse data generated from targeted sequencing platforms. Its platform-agnostic design enables seamless comparison of results across diverse sequencing technologies. CANCAN also addresses the need to evaluate alternative copy number solutions by providing transparent, quantitative likelihood estimates for each option. In addition, CANCAN offers detailed reports for potential biomarkers, highlighting both the local genomic context and the underlying raw data supporting each estimate. This level of transparency helps determine whether a variant or gene aberration represents a true biological driver in the tumor, and, importantly, informs the likelihood of a meaningful response to tailored treatment. In our bench-marking analyses, we aligned CANCAN’s results with those from multiple best-practice methods. We assessed the degree of consistency across platforms, focusing particularly on targeted sequencing data. A critical feature for methods that integrate copy number and somatic variant data is the accurate determination of tumor purity and ploidy. These parameters directly influence the copy number profile and thus the genomic interpretation context of somatic variant allele frequencies. Ideally, results from a given method would be compared against a definitive ground truth; however, no universally accepted gold standard currently exists for determining ploidy in tumor samples. To mitigate this limitation, we used the concordance of purity and ploidy estimates between CANCAN and other established methods as a key benchmarking metric in this study. Where alternative methods agreed, CANCAN demonstrated strong concordance and nearly complete alignment in the WGS and WES datasets. For benchmarking of TSO500-based results, we curated a high-confidence benchmarking dataset by leveraging matched WGS–derived estimates of tumor purity and ploidy. Notably, CANCAN achieved complete concordance between TSO500-based results and the curated WGS dataset, providing strong evidence that highly targeted sequencing data can accurately capture global genomic features such as ploidy and purity. Furthermore, CANCAN demonstrated superior concordance with WGS-derived estimates compared with PureCN, indicating performance that is at least comparable to, and potentially exceeding, that of PureCN. Utilizing a quantitative metric to assess differences in copy number estimates yielded results that closely mirrored those from purity and ploidy analyses. This was expected given the strong interdependence between these parameters and allele-specific absolute copy number estimates. For WGS data, the greatest difference in copy number estimates according to the d-metric occurred between ASCAT and PURPLE. This is in line with CANCAN demonstrating performance comparable to these established methods. Analogous analyses of WES and TSO500 data were consistent with the WGS results.

Although validation of CANCAN-derived ploidy estimates using IOD was possible for only five samples, the results provide support for the robustness of CANCAN. Three of the five samples showed strong concordance between CANCAN’s primary ploidy estimates and those obtained by IOD, consistent with the results from PureCN. In the fourth case, concordant results were achieved when considering CANCAN’s second-ranked ploidy estimate; the top-ranked solution corresponded to a doubling of the ploidy inferred from IOD. Distinguishing between such ploidy alternatives is a recognized challenge in copy number analysis, as a whole-genome duplication can produce highly similar profiles^28^. Importantly, an incorrect distinction between duplicated and non-duplicated states typically does not affect key outcomes, such as the identification of regions with loss of heterozygosity (LOH), and is therefore unlikely to meaningfully influence clinical interpretation. In this case, PureCN’s second-ranked solution did not align with the IOD estimate. The fifth sample did not yield concordant ploidy estimates for either the top-or second-ranked CANCAN solutions, and PureCN likewise failed to produce concordant results. WGS-based CANCAN analysis of an independent biopsy from the same patient indicated extensive subclonality, a condition known to violate key assumptions of copy number estimation from bulk DNA, irrespective of sequencing platform. Moreover, CANCAN’s purity estimate for this sample was low, further complicating accurate copy number inference. Together, these factors provide a reasonable explanation for the discordant results and highlight the inherent limitations of bulk DNA–based copy number analysis in samples with pronounced subclonality^29^. No evidence of inferior performance for CANCAN relative to PureCN was observed in these analyses.

## 4.1 Software availability and installation

Software with detailed instructions is available as an R package at the web site https://github.com/arnevpladsen/CANCAN. The software is open source and may be used according to the Apache License, version 2.0.

## Acknowledgments

The authors thank all the patients who have contributed to this study by donating tumor tissue and blood. The authors also thank the IMPRESS-Norway consortium for providing access to sequencing data and tumor tissue. The authors thank the the Genomics Core Facility at Oslo University Hospital, facilitating sequencing of tumor samples in the IMPRESS-Norway study. The results published here are in part based upon data generated by the TCGA Research Network: https://www.cancer.gov/tcga.

## Author contributions

A.V.P., H.G.R., and O.C.L. developed the CANCAN method with input from D.N., T.L., S.Z., S.N., O.E., and E.H. A.V.P., D.V., and O.C.L. performed the statistical and bioinformatic analyses with support from S.N., S.Z., and D.N. T.L., B.K.D., and C.W. designed and performed laboratory experiments and sequencing of IMPRESS-Norway samples. W.K. conducted the integrated ploidy analyses. G.O.H. represented the IMPRESS-Norway consortium and facilitated the infrastructure for sample collection and sequencing. A.V.P., H.G.R., and O.C.L. wrote the manuscript. All authors critically revised the manuscript and approved the final version.

## Data availability

TCGA data used in this study are publicly available from the Genomic Data Commons (GDC) portal. Sequencing data from the IMPRESS-Norway cohort are not publicly available due to patient privacy regulations but can be made available upon reasonable request and subject to appropriate ethical approvals.

