## Supplementary material for "CANCAN: high-resolution copy number and mutation heterogeneity analysis of DNA sequence data for clinical applications"

### Supplementary material and methods

#### The Cancer Genome Atlas Program (TCGA)

DNA sequencing data were obtained from a subset of 100 breast cancer patients from the TCGA cohort[1]. Aligned whole genome sequencing BAM files were downloaded from the Protected Data Cloud portal (https://bionimbus-pdc.opensciencedatacloud.org) under controlled access; the samples were part of the PCAWG consortium[2]. The whole genome sequencing data were aligned to the GRCh37 reference genome, using BWA-MEM. DNA whole exome libraries were constructed using the SeqCap EZ Exome Library v2.0 kit and sequenced on the Illumina Hi-Seq 2000 machine. At least 70% coverage at 20x depth were required to pass quality control. Aligned whole exome sequencing BAM files were downloaded from the GDC website (https://portal.gdc.cancer.gov); alignment to the GRCh38 reference genome was performed using BWA with Mark Duplicates and BQSR. Somatic variant data from WES was obtained using Mutect2[3] and was downloaded from the UCSC Xena Browser portal (https://xenabrowser.net).

#### CANCAN

TSO500 data analysis, genomic windows for read count collection were defined based on platform-specific targets. These windows were extended by up to 250 bp on each side of the target regions, depending on quality metrics from a corresponding PoN of 102 samples. Specifically, the extension was performed iteratively in 1 bp increments for each target region until fewer than 70% of the samples in the PoN provided less than 50$\times$ coverage at a given position or the maximum extension of 250 bp was reached. Neighboring genomic windows with an overlap were merged. Once the final padding was determined, windows exceeding 400 bp were iteratively split into 400 bp windows, ensuring that the final window remained larger than 300 bp.

### Supplementary Figures

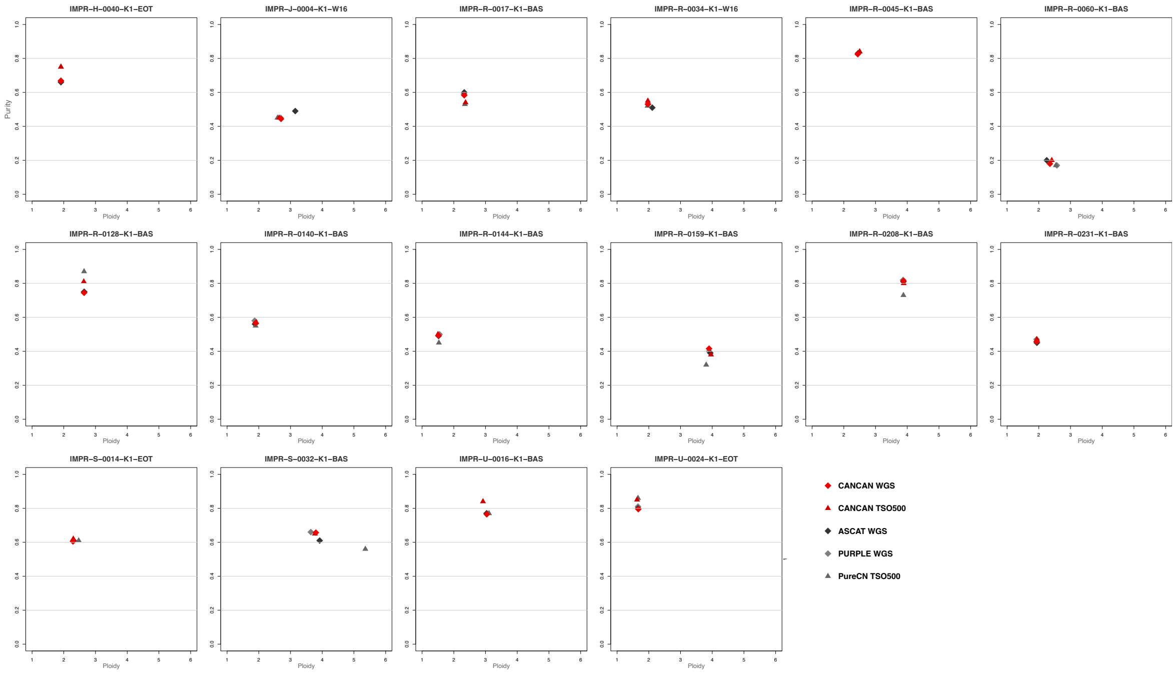


**Supplementary Figure 1** Purity and ploidy estimates for TSO500 and WGS data, stratified by copy number method.

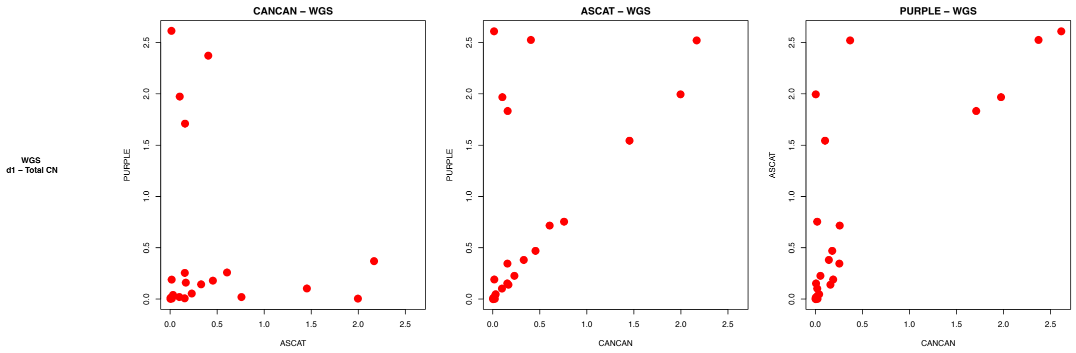


**Supplementary Figure 2** Differences in WGS-based copy number estimates according to the metric $d$, stratified by copy number method.

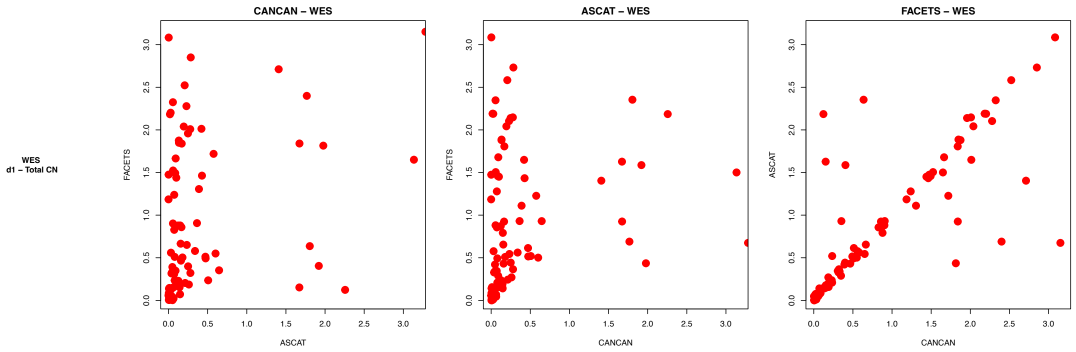


**Supplementary Figure 3** Differences in WES-based copy number estimates according to the metric $d$, stratified by copy number method.
